# Suppression of endothelial miR-22-3p mediates non-small cell lung cancer cell-induced angiogenesis

**DOI:** 10.1101/2021.04.04.438401

**Authors:** Yuan Gu, Gianni Pais, Vivien Becker, Christina Körbel, Emmanuel Ampofo, Elke Ebert, Johannes Hohneck, Nicole Ludwig, Eckart Meese, Rainer M. Bohle, Yingjun Zhao, Michael D. Menger, Matthias W. Laschke

## Abstract

MicroRNAs (miRNAs) expressed in endothelial cells (ECs) are powerful regulators of angiogenesis, which is essential for tumor growth and metastasis. Here, we demonstrated that miR-22-3p (miR-22) is preferentially and highly expressed in ECs, while its endothelial level is significantly down-regulated in human non-small cell lung cancer (NSCLC) tissues when compared to matched non-tumor lung tissues. This reduction of endothelial miR-22 is induced by NSCLC cell-secreted tumor necrosis factor (TNF)-α and interleukin (IL)-1β. Endothelial miR-22 functions as a potent angiogenesis inhibitor that inhibits all the key angiogenic activities of ECs and consequently NSCLC growth through directly targeting sirtuin (*SIRT*) *1* and fibroblast growth factor receptor (*FGFR*) *1* in ECs, leading to inactivation of AKT/mammalian target of rapamycin (mTOR) signaling. These novel findings provide insight into the molecular mechanisms of NSCLC angiogenesis and indicate that endothelial miR-22 represents a potential target for the future anti-angiogenic treatment of NSCLC.

## Introduction

Angiogenesis, i.e. the formation of new blood vessels from pre-existing ones, is essential for tumor growth and metastasis. Accordingly, excessive angiogenesis is a poor prognostic indicator for the aggressiveness of different cancer types, such as non-small cell lung cancer (NSCLC) (1). Tumor angiogenesis is tightly regulated by the balance between pro- and anti-angiogenic factors, which involves the dynamic communication between tumor cells and endothelial cells (ECs). Tumor cells are capable of releasing different pro-angiogenic factors, such as vascular endothelial growth factor (VEGF), basic fibroblast growth factor (bFGF, FGF2), epidermal growth factor (EGF), tumor necrosis factor (TNF)-α, interleukin (IL)-1β, IL-6 and IL-8 (2, 3). The binding of these factors to their receptors located on ECs activates pivotal downstream angiogenesis-related signaling pathways, such as phosphoinositide 3 kinase (PI3K)/AKT/mammalian target of rapamycin (mTOR) signaling (4). Consequently, ECs are stimulated to degrade their basement membrane, proliferate, migrate toward tumor cells and interconnect with each other to form new microvascular networks (2, 4).

Previous studies have shown that sirtuin (SIRT) 1 plays a crucial role in the regulation of angiogenesis (5). SIRT1 is a prototype member of the sirtuin family of nicotinamide adenine dinucleotide-dependent class III histone deacetylases. Loss of SIRT1 results in a significant reduction of EC sprouting and branching activity (5). Moreover, endothelial SIRT1 deletion impairs angiogenesis within ischemic hindlimbs and the kidney (5, 6). The pro-angiogenic effect of SIRT1 is most probably mediated by some of its substrates. In fact, it has been reported that SIRT1 deacetylates AKT, which binds to phosphatidylinositol (3,4,5)-triphosphate, leading to the activation of the AKT/mTOR pathway (7). In addition, SIRT1 deacetylates the forkhead transcription factor FOXO1 and, thus, suppresses its anti-angiogenic activity (5). Besides, SIRT1 can also promote the phosphorylation of AKT by up-regulating the transcription of Rictor, a component of mechanistic target of rapamycin complex 2 (mTORC2) (8).

MicroRNAs (miRNAs) are short (∼22 nucleotides), endogenous, non-coding RNAs that modulate gene expression primarily through binding to the 3’-untranslated region (UTR) of messenger RNA (mRNA), leading to mRNA degradation or translation inhibition (9). In the last decade, accumulating evidence has suggested miRNAs as powerful regulators of angiogenesis. Furthermore, miRNA deregulation has been linked to tumor development and progression. Of interest, alterations of miR-22-3p (miR-22) expression within different human body fluids and tumor tissues are considered to be of great significance for the diagnosis, surveillance and prognosis of multiple types of cancer, such as NSCLC (10). MiR-22, which is located on chromosome 17p13 and highly conserved among metazoans (11), has been reported to be also expressed in different types of ECs (12). However, its role in regulating tumor angiogenesis remains elusive.

In the present study, we analyzed the regulation of endothelial miR-22 by NSCLC cells. We then systematically investigated the function of miR-22 in basic angiogenic processes, including EC proliferation, migration and tube formation. The observed anti-angiogenic action of miR-22 was further confirmed in an *ex vivo* mouse aortic ring assay and an *in vivo* Matrigel plug assay. In addition, we studied the effects of endothelial miR-22 on tumor angiogenesis and growth in a mouse flank tumor model. Finally, mechanistic analyses identified *SIRT1* and *FGFR1* as functional targets of miR-22 in ECs.

## Results

### Endothelial miR-22 is down-regulated in human NSCLC tissues

In a first step, ECs lining the blood vessels in tumor tissues and matched adjacent non-tumor lung tissues from 12 patients with lung adenocarcinoma were retrieved by means of laser capture microdissection (LCM). By a small-scale screening using real-time PCR we identified miR-22 to be significantly down-regulated in ECs isolated from tumor tissues when compared to those isolated from matched non-tumor lung tissues (Figure 1A).

**Figure 1.**
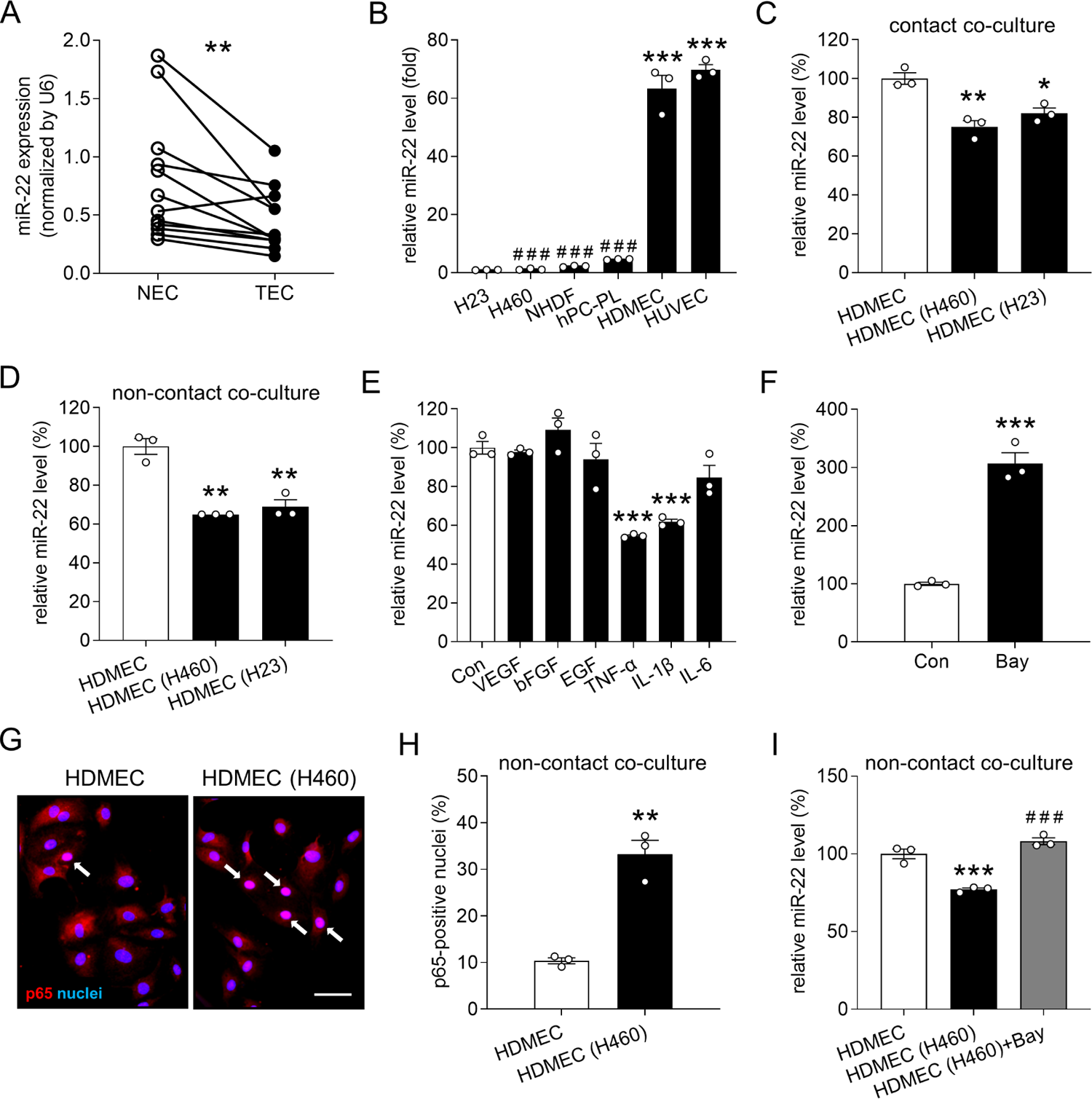
NSCLC cells down-regulate miR-22 expression in ECs. **A**: Expression level of miR-22 (normalized by U6) in ECs dissected from non-tumor (NEC) or tumor tissues (TEC) of NSCLC patients by means of LCM, as assessed by real-time PCR (n = 12). **B**: Expression level of miR-22 (in fold of H23) in NCI-H23 cells, NCI-H460 cells, NHDFs, hPC-PLs, HDMECs and HUVECs, as assessed by real-time PCR (n = 3). **C**: Expression level of miR-22 (in % of HDMEC) in isolated HDMECs that were cultured alone (HDMEC) or co-cultured in direct contact with NCI-H460 cells (HDMEC (H460)) or NCI-H23 cells (HDMEC (H23)), as assessed by real-time PCR (n = 3). **D**: Expression level of miR-22 (in % of HDMEC) in HDMECs that were cultured alone (HDMEC) or co-cultured with NCI-H460 cells (HDMEC (H460)) or NCI-H23 cells (HDMEC (H23)) without contact in a transwell plate, as assessed by real-time PCR (n = 3). **E**: Expression level of miR-22 (in % of Con) in HDMECs that were exposed for 24 h to vehicle (Con), 50 ng/mL VEGF, 50 ng/mL bFGF, 100 ng/mL EGF, 10 ng/mL TNF-α, 2 ng/mL IL-1β or 100 ng/mL IL-6 in EBM, as assessed by real-time PCR (n = 3). **F**: Expression level of miR-22 (in % of Con) in HDMECs that were treated for 24 h with vehicle (Con) or 1 µM Bay 11-7082 (Bay), as assessed by real-time PCR (n = 3). **G**: Cellular localization of NF-κB in HDMECs that were cultured alone or co-cultured with NCI-H460 cells without contact in a transwell plate and stained for p65 (red). Cell nuclei were labeled with Hoechst 33342 (blue). The nuclear translocation of p65 is indicated by arrows. Scale bar: 60 µm. **H**: p65-positive nuclei (in % of the total number of nuclei) of HDMECs that were cultured alone (HDMEC) or co-cultured with NCI-H460 cells (HDMEC (H460)) without contact in a transwell plate (n = 3). **I**: Expression level of miR-22 (in % of HDMEC) in HDMECs that were cultured alone (HDMEC) or co-cultured with NCI-H460 cells (HDMEC (H460)) without contact in a transwell plate in the absence or presence of Bay, as assessed by real-time PCR (n = 3). Means ± SEM. *P < 0.05, **P < 0.01, ***P < 0.001 vs. NEC, H23, HDMEC or Con; ^###^P < 0.001 vs. HDMEC or HDMEC (H460).

Of note, miR-22 was found to be preferentially and highly expressed in both types of analyzed ECs, i.e. human dermal microvascular endothelial cells (HDMECs) and human umbilical vein endothelial cells (HUVECs), when compared to NSCLC cells (NCI-H460 and NCI-H23) and other cell types in the tumor microenvironment, such as pericytes (human pericytes from placenta (hPC-PLs)) and fibroblasts (normal human dermal fibroblasts (NHDFs)). This indicates a specific and important regulatory function of miR-22 in ECs (Figure1B).

### NSCLC cells down-regulate miR-22 expression in ECs

Since tumor cells are capable of stimulating the angiogenic activity of ECs by both direct cell-cell contact and paracrine signaling, we next utilized a contact co-culture system to investigate how the expression of miR-22 in HDMECs is regulated by NSCLC cells. After 24 h of either culturing HDMECs alone or co-culturing them with NCI-H460 or NCI-H23 cells, HDMECs were isolated using CD31 magnetic beads. The purity of isolated HDMECs was approximately 99% and 90% in the HDMECs mono-culture and co-culture group, respectively, as assessed by flow cytometry. Real-time PCR assays revealed a 25% and a 18% reduction of miR-22 expression in HDMECs co-cultured with NCI-H460 cells and NCI-H23 cells, when compared to HDMEC mono-culture (Figure 1C). In an additional set of experiments, we co-cultured HDMECs with NSCLC cells, however, without contact between these two cell types in a transwell plate. Interestingly, this non-contact co-culture with NCI-H460 cells caused a 35% decrease in the miR-22 expression level of HDMECs (Figure 1D), indicating that soluble factors secreted by the tumor cells contribute to the down-regulation of endothelial miR-22. This finding was confirmed by the co-culture of HDMECs with NCI-H23 cells, which also significantly reduced the endothelial expression of miR-22 by 31% (Figure 1D).

In order to identify individual factors mediating the NSCLC cell-induced reduction of endothelial miR-22, HDMECs were stimulated with the growth factors VEGF, bFGF and EGF as well as the pro-inflammatory cytokines TNF-α, IL-1β and IL-6. Real-time PCR analyses revealed that the expression of miR-22 is significantly suppressed by TNF-α and IL-1β, but not affected by VEGF, bFGF, EGF and IL-6 stimulation (Figure 1E). Given the fact that both TNF-α and IL-1β are upstream inducers of nuclear factor (NF)-κB, which promotes or represses the transcription of a broad spectrum of genes and miRNAs (13, 14), we then investigated whether NF-κB inhibits the transcription of miR-22 in ECs. For this purpose, HDMECs were exposed to the NF-κB inhibitor Bay 11-7082 (Bay) for 24 h. This resulted in a 2-fold increase of miR-22 expression when compared to vehicle-treated controls (Figure 1F), indicating that this miRNA is transcriptionally repressed by NF-κB.

To investigate whether NF-κB mediates the down-regulation of endothelial miR-22 induced by NSCLC cells, we assessed the activation status of NF-κB in HDMECs cultured alone or co-cultured with NCI-H460 cells without contact. By means of immunofluorescence, we demonstrated that the nuclear translocation of p65, a main subunit of NF-κB, is significantly enhanced in HDMECs co-cultured with tumor cells (Figure 1G and H). Importantly, blockade of NF-κB signaling with Bay completely reversed the reduction of endothelial miR-22 induced by non-contact co-culture with NCI-H460 cells (Figure 1I).

### MiR-22 inhibits the angiogenic activity of ECs

To study the function of miR-22 in regulating EC angiogenic activity, we transfected HDMECs with miR-22 mimic (miR-22m) and miR-22 inhibitor (miR-22i) to up- and down-regulate the intracellular level of this miRNA, respectively. Cells transfected with negative control of mimic (NCm) or negative control of inhibitor (NCi) served as controls. The transfection efficiencies of miR-22m (5 nM) and miR-22i (100 nM) were evaluated by real-time PCR assays, as shown in Figure 2-figure supplement 1A and B.

**Figure 2.**
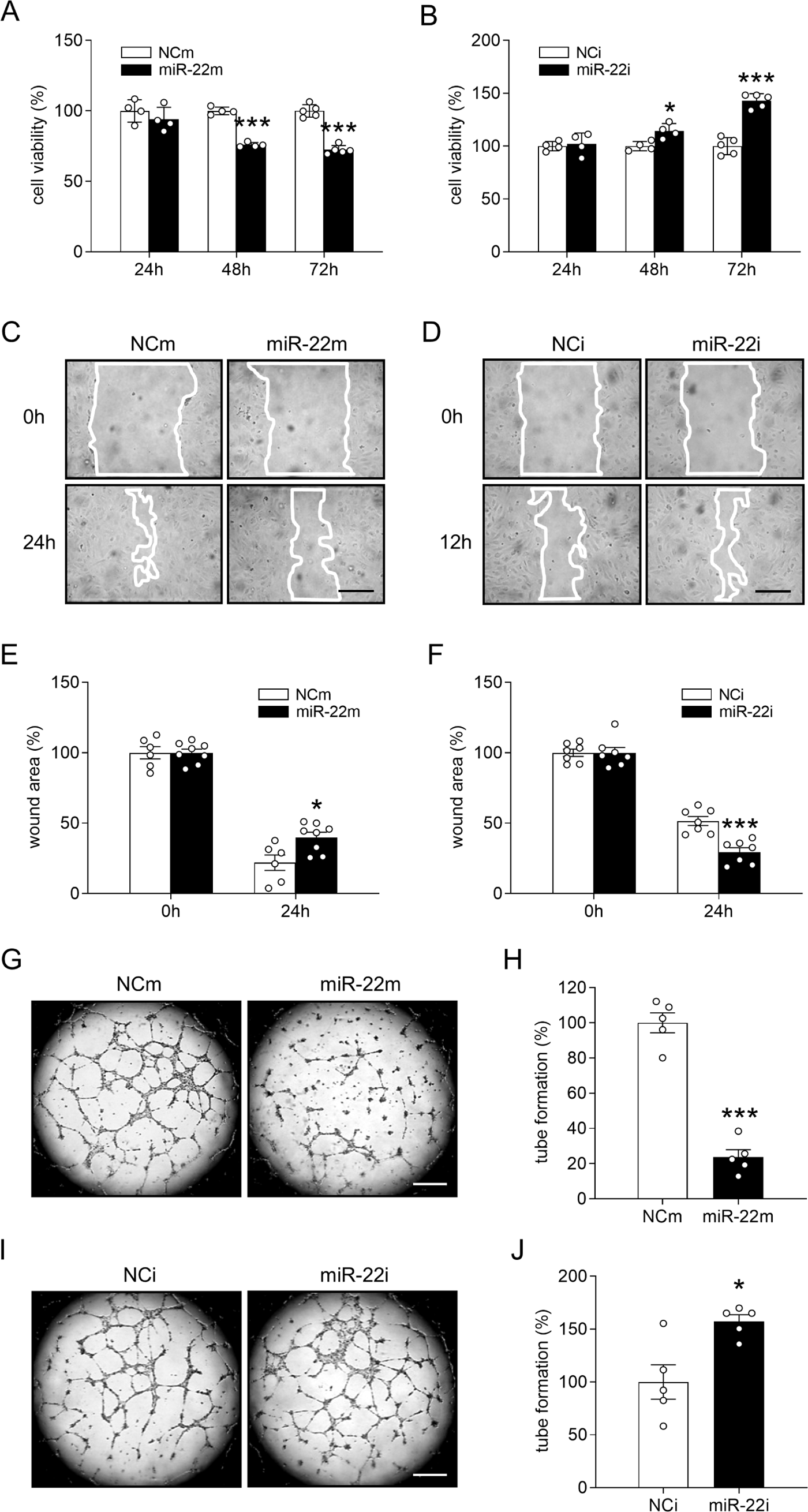
MiR-22 inhibits HDMEC viability, migration and tube formation. **A**, **B**: Viability (in % of NCm or NCi) of HDMECs transfected with miR-22m (**A**), miR-22i (**B**) or corresponding scrambled NCm (**A**) and NCi (**B**), as assessed by WST-1 assay (n = 4-5). After transfection, the cells were reseeded in 96-well plates and cultured for 24 h, 48 h or 72 h. **C**, **D**: Phase-contrast microscopic images of HDMECs at 0 h, 12 h or 24 h after scratching. The cells were transfected with miR-22m (**C**), miR-22i (**D**) or corresponding scrambled NCm (**C**) and NCi (**D**). White lines indicate scratched wound area. Scale bars: 190 µm. **E**, **F**: Wound area (in % of 0 h) created by scratching the monolayer of HDMECs transfected with miR-22m (**E**), miR-22i (**F**) or corresponding scrambled NCm (**E**) and NCi (**F**), as assessed by scratch wound healing assay (n = 6-8). **G**, **I**: Phase-contrast microscopic images of tube-forming HDMECs. The cells were transfected with miR-22m (**G**), miR-22i (**I**) or corresponding scrambled NCm (**G**) and NCi (**I**). Scale bars: 550 µm. **H**, **J**: Tube formation (in % of NCm or NCi) of HDMECs transfected with miR-22m (**H**), miR-22i (**J**) or corresponding scrambled NCm (**H**) and NCi (**J**), as assessed by tube formation assay (n = 5). Means ± SEM. *P < 0.05, **P < 0.01, ***P < 0.001 vs. NCm or NCi.

At first, water-soluble tetrazolium (WST)-1 assays were performed to assess the viability of ECs. Transfection with miR-22m significantly reduced the viability of HDMECs after 48 h of incubation (Figure 2A). This inhibitory effect of miR-22m was detectable for at least 10 days (Figure 2-figure supplement 1C). In contrast, an increased viability rate was observed in miR-22i-transfected ECs (Figure 2B). The effect of miR-22 on EC proliferation was further analyzed by flow cytometry assessing the cell cycle distribution of transfected HDMECs. The S-phase cell population was significantly increased in miR-22m-transfected HDMECs when compared to NCm-transfected controls (Figure 2-figure supplement 2A and B). This was associated with an increase in the number of sub-G1-phase cells (Figure 2-figure supplement 2A and C). These results suggest that miR-22 inhibits EC proliferation and induces apoptosis by blocking the cells in the S phase.

To investigate the function of miR-22 in regulating EC motility, scratch wound healing assays and transwell migration assays were performed. Transfection of HDMECs with miR-22m markedly delayed the healing of scratched wounds (Figure 2C and E) and reduced the number of transwell migrated cells by 34% (Figure 2-figure supplement 3A and B). In contrast, transfection of HDMECs with miR-22i significantly promoted wound closure (Figure 2D and F) and enhanced cell migration by 42% (Figure 2-figure supplement 3C and D).

In addition, we performed a tube formation assay to investigate the function of miR-22 in regulating the tube forming activity of HDMECs. Transfection with miR-22m markedly reduced the number of newly developed tube meshes by 76% when compared to NCm-transfected controls (Figure 2G and H). In contrast, miR-22i significantly augmented EC tube formation by 64% (Figure 2I and J).

### Endothelial miR-22 suppresses angiogenesis *ex vivo* and *in vivo*

To elucidate whether miR-22 is involved in endothelial sprouting, we performed an *ex vivo* mouse aortic ring assay. We found that the area of vascular sprouting from aortic rings is significantly decreased by transfection with miR-22m (Figure 3A and B) and significantly increased by transfection with miR-22i (Figure 3C and D).

**Figure 3.**
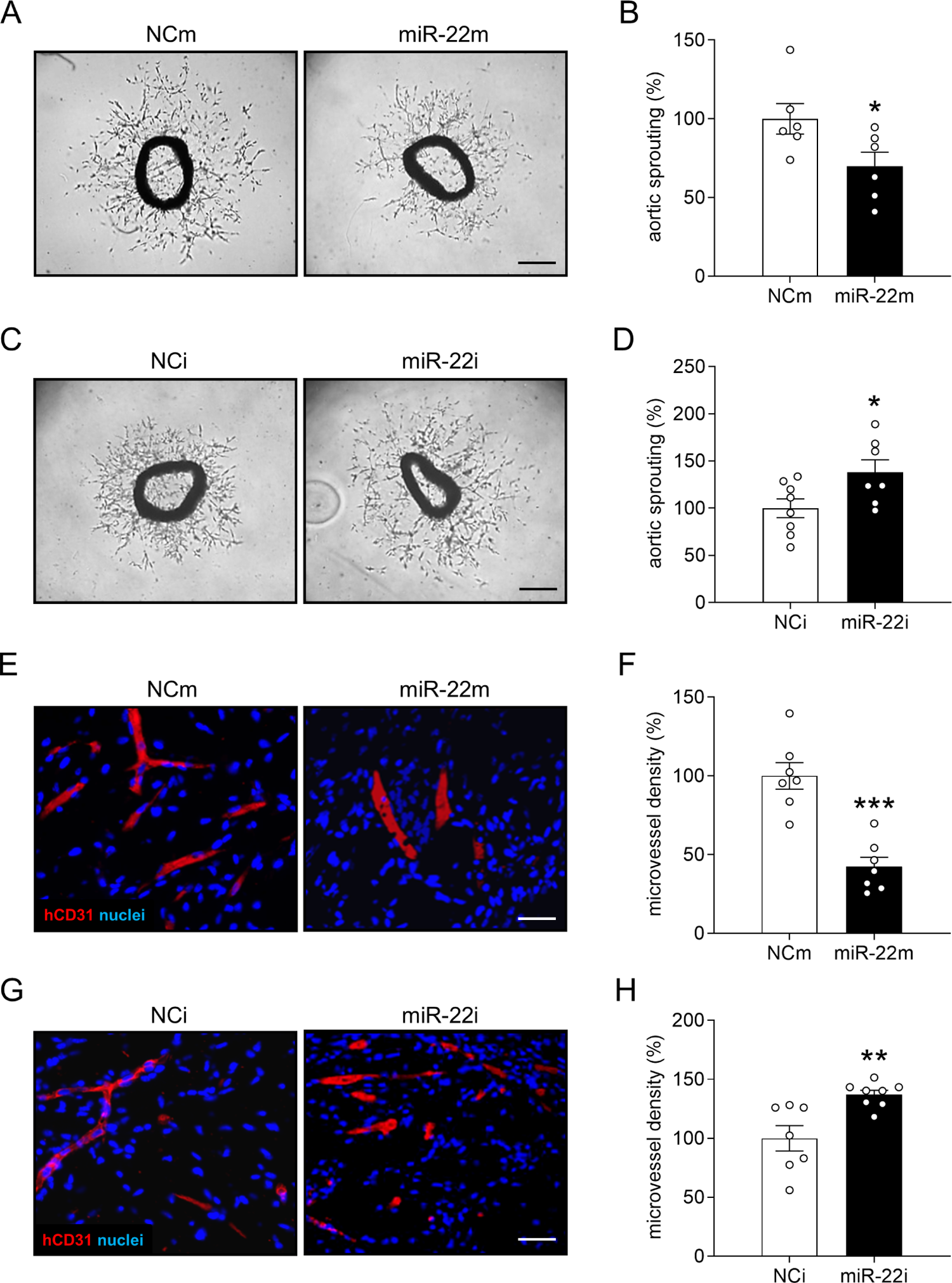
MiR-22 suppresses angiogenesis ex vivo and in vivo. **A**, **C**: Phase-contrast microscopic images of mouse aortic rings, which were transfected with miR-22m (**A**), miR-22i (**C**) or corresponding scrambled NCm (**A**) and NCi (**C**) overnight and then cultured in Matrigel for 6 days. Scale bars: 400 µm. **B**, **D**: Sprouting (in % of NCm or NCi) of aortic rings that were transfected with miR-22m (**B**), miR-22i (**D**) or corresponding scrambled NCm (**B**) and NCi (**D**), as assessed by computer-assisted image analysis (n = 6-8). **E**, **G**: Immunohistochemical detection of human CD31-positive microvessels (red) in Matrigel plugs containing HDMECs transfected with miR-22m (**E**), miR-22i (**G**) or corresponding scrambled NCm (**E**) and NCi (**G**). Sections were additionally stained with Hoechst 33342 to identify cell nuclei (blue). Scale bars: 40 µm. **F**, **H**: Microvessel density (in % of NCm or NCi) of Matrigel plugs containing HDMECs transfected with miR-22m (**F**), miR-22i (**H**) or corresponding scrambled NCm (**F**) and NCi (**H**), as assessed by immunohistochemistry (n = 7-8). Means ± SEM. *P<0.05, **P < 0.01, ***P < 0.001 vs. NCm or NCi.

To confirm our *in vitro* findings, we performed an *in vivo* Matrigel plug assay. Matrigel plugs containing miR-22m-transfected HDMECs exhibited a 58% reduction of the microvessel density 7 days after implantation when compared to those containing NCm-transfected controls (Figure 3E and F). In contrast, plugs containing miR-22i-transfected cells presented with a 42% higher microvessel density than plugs containing NCi-transfected cells (Figure 3G and H).

### Endothelial miR-22 inhibits tumor angiogenesis and growth

The findings above demonstrated that: i) NSCLC cells down-regulate the expression level of miR-22 in ECs and ii) miR-22 acts as a potent angiogenesis inhibitor. Hence, we assumed that tumor cells stimulate angiogenesis at least partially through suppressing endothelial miR-22 expression. To verify this hypothesis, we established an *in vivo* tumor cell-EC communication model by injecting NCI-H460 cells together with NCm- or miR-22m-transfected HDMECs into the flanks of NOD-SCID mice. Digital caliper measurements and high-resolution ultrasound imaging were performed to assess the volume of the newly developing tumors. We found that transfection of HDMECs with miR-22m significantly inhibits NCI-H460 tumor development between day 7 to 14 when compared to NCm-transfected controls (Figure 4A, C and D). Accordingly, tumors containing miR-22m-transfected HDMECs also exhibited a markedly reduced final tumor weight (Figure 4B). As expected, overexpression of miR-22 in HDMECs significantly counteracted the tumor cell-stimulated development of human microvessels within the tumors, but not the angiogenic ingrowth of mouse microvessels from the surrounding host tissue (Figure 4E and F). Additional immunohistochemical analyses demonstrated that tumors containing miR-22m-transfected HDMECs exhibited less Ki67-positive but more cleaved caspase (casp)-3-positive tumor cells when compared to controls (Figure 4G-J). This indicates that miR-22 overexpression in tumor ECs inhibits the proliferation of tumor cells and also promotes their apoptotic cell death.

**Figure 4.**
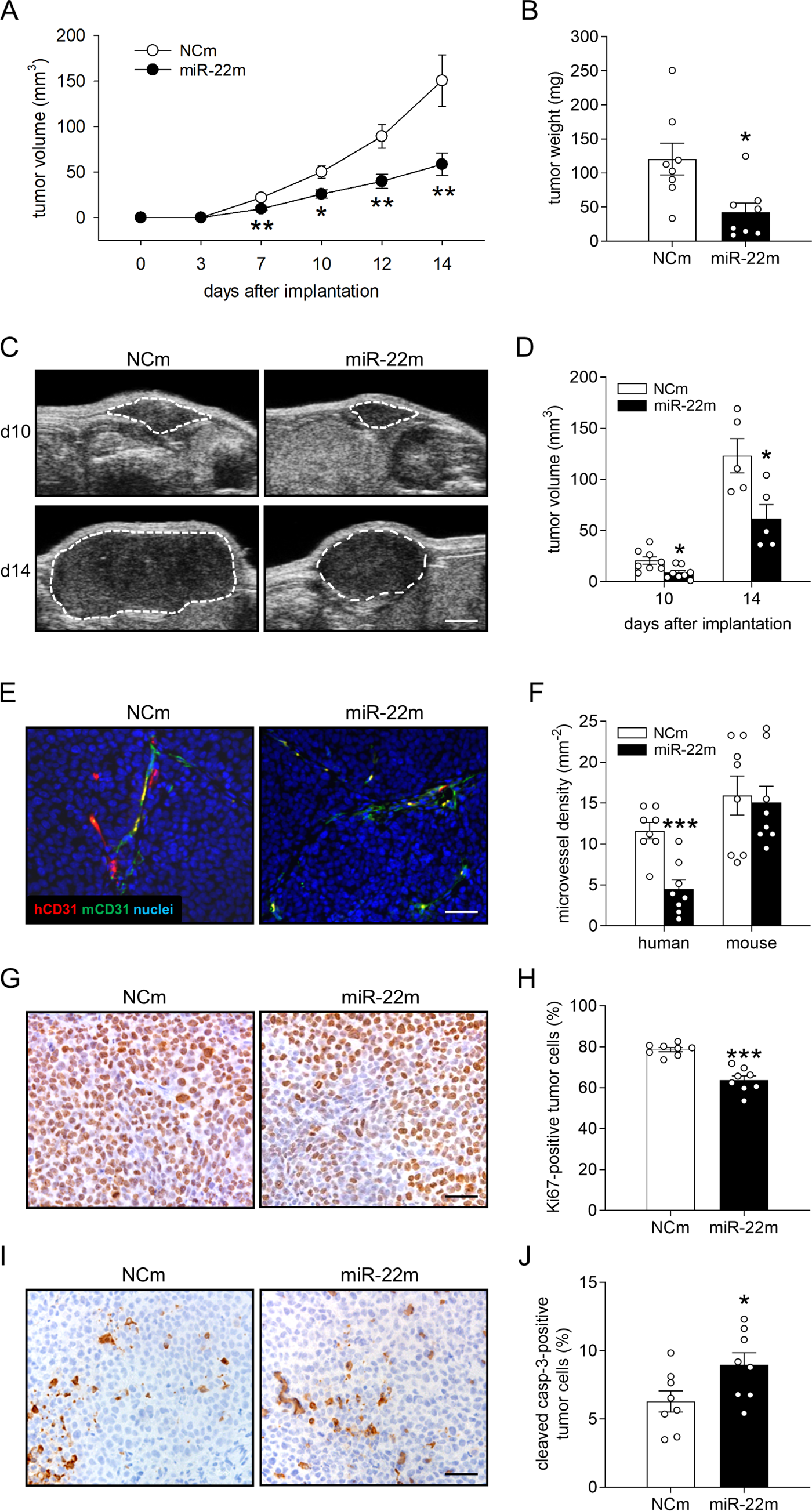
Endothelial miR-22 inhibits tumor angiogenesis and growth. **A**: Volume (mm³) of developing NCI-H460 flank tumors containing NCm- or miR-22m-transfected HDMECs, as assessed by means of a digital caliper on the day of tumor induction (day 0) as well as on day 3, 7, 10, 12 and 14 (n = 8). **B**: Final weight (mg) of tumors containing NCm- or miR-22m-transfected HDMECs on day 14 (n = 8). **C**: High-resolution ultrasound imaging of tumors containing NCm- or miR-22m-transfected HDMECs on day 10 and 14 after implantation. The borders of tumors are marked by white dashed lines. Scale bar: 1.8 mm. **D**: Volume (mm³) of tumors containing NCm- or miR-22m-transfected HDMECs, as assessed by high-resolution ultrasound imaging on day 10 and 14 (n = 5-8). **E**: Immunohistochemical detection of newly formed human (red) and mouse (green) microvessels in tumors containing NCm- or miR-22m-transfected HDMECs on day 14 (n = 8). Sections were stained with Hoechst 33342 to identify cell nuclei (blue). Scale bar: 60 µm. **F**: Density (mm^-2^) of human and mouse microvessels in tumors containing NCm- or miR-22m-transfected HDMECs on day 14 (n = 8). **G**, **I**: Immunohistochemical detection of human Ki67-(**G**) or cleaved casp-3-positive (**I**) tumor cells within NCI-H460 xenografts containing NCm- or miR-22m-transfected HDMECs. Scale bars: 25 µm. **H**, **J**: Ki67-positive (**H**) or cleaved casp-3-positive cells (**J**) (in % of the total number of nuclei) within NCI-H460 xenografts containing NCm- or miR-22m-transfected HDMECs (n = 8). Means ± SEM. *P<0.05, **P < 0.01, ***P < 0.001 vs. NCm.

### MiR-22 targets *SIRT1* and *FGFR1* in ECs

To identify the functional targets of miR-22 that mediate its anti-angiogenic effects in ECs, we first analyzed the predicted human target genes of miR-22 according to the algorithms of miRDB and TargetScan. We detected 5 genes that are involved in angiogenesis and have not been validated as miR-22 targets, which encode tumor necrosis factor receptor (TNFR) 2, vascular endothelial zinc finger (VEZF) 1, transforming growth factor beta-activated kinase (TAK) 1, serine–arginine protein kinase (SRPK) 1 and protein kinase C beta (PRKCB). However, none of these genes was down-regulated in miR-22m-transfected HDMECs when compared to NCm-transfected controls (Figure 5A).

**Figure 5.**
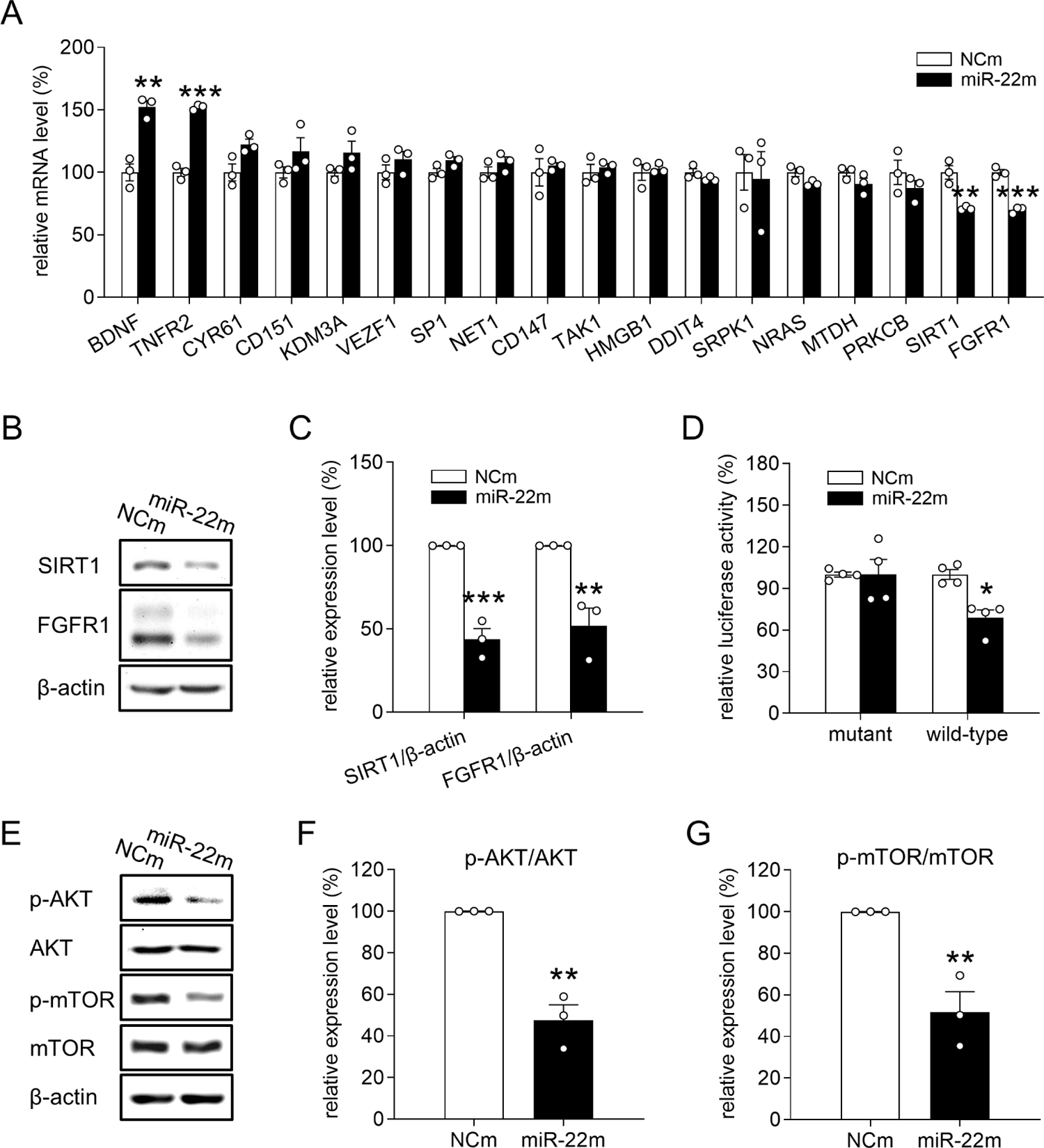
MiR-22 targets *SIRT1* and *FGFR1* in ECs. **A**: mRNA levels (in % of NCm) of putative and validated human target genes of miR-22 in NCm- or miR-22m-transfected HDMECs, as assessed by real-time PCR (n = 3). **B**: Western blot of SIRT1, FGFR1 and β-actin expression in HDMECs transfected with NCm or miR-22m. **C**: Expression level (in % of NCm) of SIRT1/β-actin and FGFR1/β-actin, as assessed by Western blot (n = 3). **D**: Luciferase activity (in % of NCm) in 293T cells co-transfected with NCm or miR-22m and a reporter plasmid carrying mutant or wild-type *FGFR1*-3’UTR, as assessed by luciferase assay (n = 4). **E**: Western blot of p-AKT, AKT, p-mTOR, mTOR and β-actin expression in HDMECs transfected with NCm or miR-22m. **F**, **G**: Expression levels (in % of NCm) of p-AKT/AKT (**F**) and p-mTOR/mTOR (**G**), as assessed by Western blot (n = 3). Means ± SEM. *P < 0.05, **P < 0.01, ***P < 0.001 vs. NCm.

Moreover, we analyzed the validated human targets of this miRNA based on the current literature and found 13 angiogenesis-related genes. These genes encode brain-derived neurotrophic factor (BDNF), cysteine-rich protein (CYR) 61, cluster of differentiation (CD) 151, lysine-specific demethylase (KDM) 3A, specificity protein (SP) 1, neuroepithelial cell transforming (NET) 1, CD147, high mobility group box protein (HMGB) 1, DNA damage inducible transcript (DDIT) 4, neuroblastoma RAS viral oncogene homolog (NRAS), metadherin (MTDH), SIRT1 and FGFR1. By performing real-time PCR assays, the mRNA levels of *SIRT1* and *FGFR1* were found to be significantly decreased in miR-22m-transfected HDMECs when compared to NCm-transfected controls (Figure 5A). Consistently, the protein levels of *SIRT1* and *FGFR1* were markedly decreased by miR-22 overexpression, as assessed by Western blot (Figure 5B and C). Recently, Hu et al. reported that miR-22 targets *FGFR1* in human liver Huh7 cells (15). We further confirmed this finding in 293T cells, which is a highly transfectable cell line and widely used for miRNA target validation. For this purpose, a dual luciferase assay was performed by co-transfecting miR-22m and *FGFR1*-3’UTR luciferase reporter plasmid (wild-type) or an empty plasmid with deletion of *FGFR1*-3’UTR (mutant) into the cells. We found that miR-22m significantly attenuates the activity of *FGFR1*-3’UTR luciferase reporter, whereas no reduction was detected upon co-transfection with mutant plasmid (Figure 5D).

Given the fact that both SIRT1 and FGFR1 are upstream proteins of the pivotal angiogenesis regulatory pathway AKT/mTOR, we performed Western blot analyses to assess the activation of this pathway in NCm- and miR-22m-transfected HDMECs. As expected, transfection with miR-22m markedly reduced the phosphorylation of AKT and mTOR by 52% and 48%, respectively (Figure 5E-G).

### MiR-22 inhibits angiogenesis through targeting *SIRT1* and *FGFR1*

Previous studies suggest an important role of SIRT1 and FGFR1 in regulating angiogenesis (5, 16). To determine whether miR-22 inhibits the angiogenic activity of ECs through targeting *SIRT1* and *FGFR1*, the specific SIRT1 inhibitor EX-527 (EX) and the selective FGFR1 inhibitor PD173074 (PD) were used in an additional panel of *in vitro* assays. By means of a WST-1 assay, we found that 10-50 µM EX and 50-500 nM PD significantly reduce the viability of HDMECs after 3 days of treatment (Figure 6A and B). Accordingly, to avoid cytotoxic effects of these compounds, we chose a minimal effective dose of each inhibitor, i.e. 10 µM EX and 50 nM PD, for the following WST-1, scratch wound healing and tube formation assays. These functional analyses revealed that exposure to EX and PD completely reverses miR-22i-promoted HDMEC viability, migration and tube formation (Figure 6C-E).

**Figure 6.**
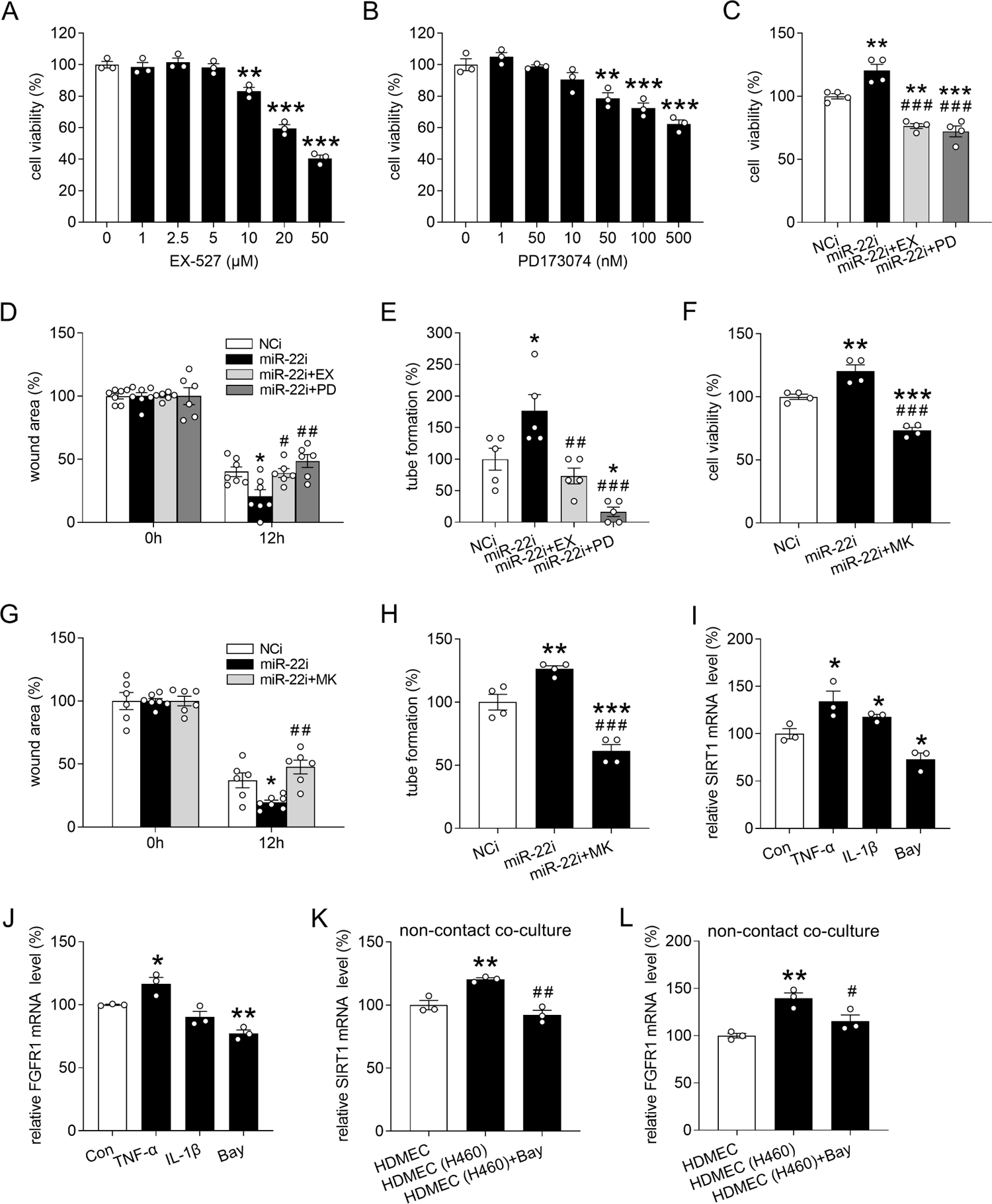
MiR-22 inhibits angiogenesis through targeting *SIRT1* and *FGFR1*. **A**, **B**: Viability (in % of 0 µM or 0 nM) of HDMECs that were exposed for 72 h to serial dilutions of EX-527 (**A**) and PD173074 (**B**), as assessed by WST-1 assay (n = 3). **C**: Viability (in % of NCi) of HDMECs that were transfected with NCi or miR-22i and then treated with 10 µM EX-527 (EX) or 50 nM PD173074 (PD) for 72 h, as assessed by WST-1 assay (n = 4). **D**: Wound area (in % of 0 h) created by scratching the monolayer of HDMECs that were transfected with NCi or miR-22i and then treated with 10 µM EX or 50 nM PD for 12 h, as assessed by scratch wound healing assay (n = 6-7). **E**: Tube formation (in % of NCi) of HDMECs that were transfected with NCi or miR-22i and then treated with 10 µM EX or 50 nM PD for 18 h, as assessed by tube formation assay (n = 5). **F**: Viability (in % of NCi) of HDMECs that were transfected with NCi or miR-22i and then treated with 5 µM MK-2206 (MK) for 72 h, as assessed by WST-1 assay (n = 4). **G**: Wound area (in % of 0 h) created by scratching the monolayer of HDMECs that were transfected with NCi or miR-22i and then treated with 5 µM MK for 12 h, as assessed by scratch wound healing assay (n = 6). **H**: Tube formation (in % of NCi) of HDMECs that were transfected with NCi or miR-22i and then treated with 5 µM MK for 18 h, as assessed by tube formation assay (n = 4). **I**, **J**: mRNA levels of *SIRT1* (**I**) and *FGFR1* (**J**) (in % of Con) in HDMECs that were exposed for 72 h to vehicle (Con), 10 ng/mL TNF-α, 2 ng/mL IL-1β or 1 µM Bay, as assessed by real-time PCR (n = 3). **K**, **L**: mRNA level of *SIRT1* (**K**) or *FGFR1* (**L**) (in % of HDMEC) in HDMECs that were cultured alone (HDMEC) or co-cultured with NCI-H460 cells (HDMEC (H460)) without contact in a transwell plate in the absence or presence of 1 µM Bay for 72 h, as assessed by real-time PCR (n = 3). Means ± SEM. *P < 0.05, **P < 0.01, ***P < 0.001 vs. 0 µM or 0 nM, NCi, Con or HDMEC; ^#^P < 0.05, ^##^P < 0.01, ^###^P < 0.001 vs. miR-22i or HDMEC (460).

Furthermore, we analyzed whether miR-22 functions through suppressing AKT/mTOR signaling, which is a common down-stream pathway of SIRT1 and FGFR1, using the highly specific AKT inhibitor MK-2206 (MK). In a previous publication (17), we found that 5-40µM MK-2206 significantly reduces HDMEC viability after 3 days of incubation. Accordingly, miR-22i-transfected HDMECs were exposed to 5 µM MK-2206 followed by WST-1, scratch wound healing and tube formation assays. By this, we could demonstrate that inhibition of AKT completely counteracts miR-22i-enhanced HDMEC viability, migration and tube formation (Figure 6F-H).

Because we found that NSCLC cells down-regulate endothelial miR-22 by activating NF-κB possibly via secreting TNF-α and IL-1β, we investigated the regulation of the miR-22 targeted genes in ECs. For this purpose, we assessed the expression of *SIRT1* and *FGFR1* in TNF-α-, IL-1β- or Bay-exposed HDMECs as well as HDMECs co-cultured with NCI-H460 cells. Real-time PCR assays revealed that TNF-α significantly increases the mRNA levels of *SIRT1* and *FGFR1* and IL-1β promotes the expression of *SIRT1* but not of *FGFR1* (Figs. 6I and J). In contrast, Bay reduced the expression of the two genes (Figure 6I and J). Moreover, non-contact co-culture of HDMEC with NCI-H460 cells significantly up-regulated the endothelial expression of *SIRT1* and *FGFR1*, whereas inhibition of NF-κB with Bay reversed this up-regulation (Figure 6K and L).

## Discussion

MiR-22 is widely studied in tumorigenesis, where it acts as a tumor suppressor or an oncogene by regulating the proliferation, migration, invasion, metastasis, apoptosis, senescence and epithelial-mesenchymal transition of different types of tumor cells (11). Moreover, the aberrant expression of miR-22 in tumor tissues and body fluids of cancer patients provides the possibility to use this miRNA as an independent diagnostic and prognostic biomarker (10). Besides, it is known that miR-22 induces endothelial progenitor cell senescence and its injection into zebrafish embryos causes defective vascular development (18, 19). However, the regulation, function and targets of miR-22 in ECs still need to be clarified. Our novel findings now demonstrate that miR-22 is preferentially and highly expressed in ECs and the suppression of endothelial miR-22 mediates NSCLC cell-promoted blood vessel formation. In fact, NSCLC cell-released TNF-α and IL-1β activate endothelial NF-κB and, thus, markedly reduce the high expression of miR-22 in ECs. This increases the angiogenic activity of ECs, because miR-22 functions as a potent angiogenesis inhibitor by targeting *SIRT1* and *FGFR1*.

Lung cancer is the most frequently diagnosed cancer and the leading cause of cancer death in both sexes worldwide (20). Non-small cell lung cancer (NSCLC), including adenocarcinoma, large cell carcinoma and squamous cell carcinoma, accounts for approximately 85% of all lung cancer cases. Despite recent advances in diagnosis and treatment, many patients with NSCLC still have limited treatment options and a poor prognosis (21). Therefore, we focused in the present study on this specific tumor type and identified miR-22 to be significantly down-regulated in ECs dissected from human NSCLC tissues when compared to that from matched non-tumor lung tissues. *In vitro*, we also detected a significantly down-regulated expression of miR-22 in HDMECs directly co-cultured with NCI-H460 or NCI-H23 cells when compared to EC mono-cultures. This is in line with a previous study reporting that miR-22 expression in primary human brain microvascular ECs is reduced by contact co-culture with U87 glioma cells (22). Hence, endothelial miR-22 seems to be regulated by different types of tumors.

Tumor cells can directly interact with ECs via adhesion receptors and gap junctions. In addition, they can activate ECs by secreting soluble factors and microvesicles into the extracellular space as well as by changing the pH, oxygen and nutrient levels in the surrounding microenvironment (23). Therefore, we next assessed the endothelial expression of miR-22 in a non-contact co-culture system, in which HDMECs and NCI-H460 or NCI-H23 cells were separated from each other in a transwell plate. In this setting, the expression of miR-22 was also significantly reduced in co-cultured HDMECs, indicating that the change in endothelial miR-22 expression is at least partially due to an indirect interaction between NSCLC cells and ECs.

To identify the factors, which mediate the communication between tumor cells and ECs, we stimulated HDMECs with several soluble factors that can be secreted by NCLSC cells and are crucially involved in angiogenesis. Our results showed that TNF-α and IL-1β, but not VEGF, bFGF, EGF and IL-6, markedly reduce the endothelial expression of miR-22. Of note, TNF-α is a major pro-inflammatory cytokine, which exerts contradictory effects on blood vessel formation. High doses of exogenous TNF-α have been shown to inhibit angiogenesis, whereas low doses or endogenous TNF-α stimulate the angiogenic process and stabilize the newly developing microvascular networks within tumors (24, 25). Moreover, Sainson et al. reported that pulsed administration of high doses of TNF-α stimulates angiogenesis by inducing a tip cell phenotype (26). In contrast, the pro-inflammatory cytokine IL-1β is widely accepted as a pro-angiogenic factor (27). TNF-α and IL-1β exert their biological functions through binding to TNF receptor and IL-1 receptor, respectively. This, in turn, recruits and activates the inhibitor of NF-kB (IkB) kinase complex. The consequent phosphorylation of IkB proteins leads to the translocation of NF-kB into the nucleus, where it promotes or represses the transcription of mRNAs and miRNAs (13, 14). Of interest, a recent study identified two NF-κB binding motifs in the miR-22 promoter that mediate the transcriptional repression of miR-22 in 182^R^-6 breast cancer cells (28). Our results now demonstrate that the exposure of HDMECs to the NF-κB inhibitor Bay 11-7082 significantly increases miR-22 expression, indicating that miR-22 is not only transcriptionally repressed by NF-κB in tumor cells but also in ECs. More importantly, we verified that NSCLC cell-induced NF-κB activation by secreting TNF-α and IL-1β contributes to the down-regulation of miR-22 in HDMECs co-cultured with NCI-H460 cells.

We next investigated the effects of endothelial miR-22 on angiogenesis. By a panel of well-established *in vitro* angiogenesis assays, we could demonstrate that miR-22 is a pleiotropic angiogenesis inhibitor that targets all the major steps of the angiogenic process, including EC proliferation, migration and tube formation. Of note, the inhibitory effects of miR-22 on these steps were not directly dependent on each other. This is indicated by the observation that miR-22m inhibits HDMEC migration and tube formation within 24 h after transfection without affecting the viability of the cells. Our *in vitro* results were further confirmed by an *ex vivo* mouse aortic ring assay and an *in vivo* Matrigel plug assay. The fact that the mouse aortic ring assay is based on the angiogenic sprouting activity of murine ECs shows that the anti-angiogenic effect of miR-22 is reproducible in ECs of different origin.

The findings above suggest that NSCLC cells stimulate angiogenesis by down-regulating endothelial miR-22 expression to support their growth. To verify this conclusion, we established an *in vivo* tumor cell-EC communication model. In this model, NCI-H460 cells admixed with NCm- or miR-22-transfected HDMECs were injected into the flanks of immunodeficient mice. By this, we could demonstrate that overexpression of miR-22 in HDMECs significantly suppresses their assembly into new microvessels within the tumors, resulting in a reduced tumor growth. Noteworthy, this modified flank tumor model only allows the manipulation of miR-22 expression in exogenous human ECs but not endogenous mouse ECs. However, these mouse ECs invade the developing tumor, assemble into new microvessels and, thus, also support tumor growth. Accordingly, our model may underestimate the inhibitory effect of miR-22 on NSCLC growth. So far, targeted delivery of miRNA into vascular ECs *in vivo* is still a big challenge (29). This largely prevents basic studies to translate into novel clinical applications. Hence, it will be necessary to develop miRNA modifications and sophisticated delivery systems to improve the safety, efficiency and specificity of miRNA-based therapeutics. Rapid progress in chemical and bioengineering of miRNA, nanotechnology and viral vector development may markedly contribute to achieve this in the future.

MiRNAs have the potential to regulate multiple target genes and related manifold signaling pathways. Moreover, each miRNA may function differently in diverse cell types due to the high complexity of cellular physiology (30). Therefore, it was necessary in the present study to identify the specific functional targets of miR-22 in ECs. For this purpose, we analyzed the putative and validated human target genes of miR-22 and identified *SIRT1* and *FGFR1* to be down-regulated in miR-22-overexpressing HDMECs. FGFR1, a member of FGFR family of receptor tyrosine kinases, is most commonly expressed on ECs (16). Activation of FGFR1 by heparin-binding FGFs, mainly FGF1 and bFGF, increases the angiogenic activity of ECs *in vitro* and *in vivo* (16). Thus, FGFR1 has been increasingly considered to be an attractive target for the anti-angiogenic treatment of tumors. In order to investigate whether the suppression of *SIRT1* or *FGFR1* mediates the anti-angiogenic function of miR-22, we exposed miR-22i-transfected HDMECs to the SIRT1 inhibitor EX-527 or the FGFR1 inhibitor PD173074. These small molecular inhibitors were used instead of short interfering RNAs (siRNAs) against SIRT1 or FGFR1, because we found in preliminary experiments that the co-transfection efficiency of miR-22i and siRNAs is quite low in HDMECs. Our results showed that both EX-527 and PD173074 completely reverse miR-22i-induced HDMEC proliferation, migration and tube formation. Hence, *SIRT1* and *FGFR1* are functional targets of miR-22 in the regulation of angiogenesis. Moreover, we found that the endothelial expression of *SIRT1* and *FGFR1* is up-regulated by NSCLC cell-activated NF-κB signaling possibly via secretion of TNF-α and IL-1β.

In conclusion, this study demonstrates that down-regulation of endothelial miR-22 significantly contributes to NSCLC cell-stimulated angiogenesis. As summarized in Figure 7, tumor cell-released TNF-α and IL-1β bind to their receptors located on ECs, causing the intracellular activation of NF-κB. This, in turn, suppresses endothelial miR-22 expression. MiR-22 targets the two pivotal pro-angiogenic regulators *SIRT1* and *FGFR1*, which results in the blockage of AKT/mTOR signaling and inhibition of angiogenesis. Thus, the NF-κB-induced suppression of miR-22 results in an increased SIRT1- and FGFR1-mediated angiogenesis. Taken together, this novel mechanism indicates that endothelial miR-22 may represent a promising therapeutic target for the treatment of NSCLC.

**Figure 7.**
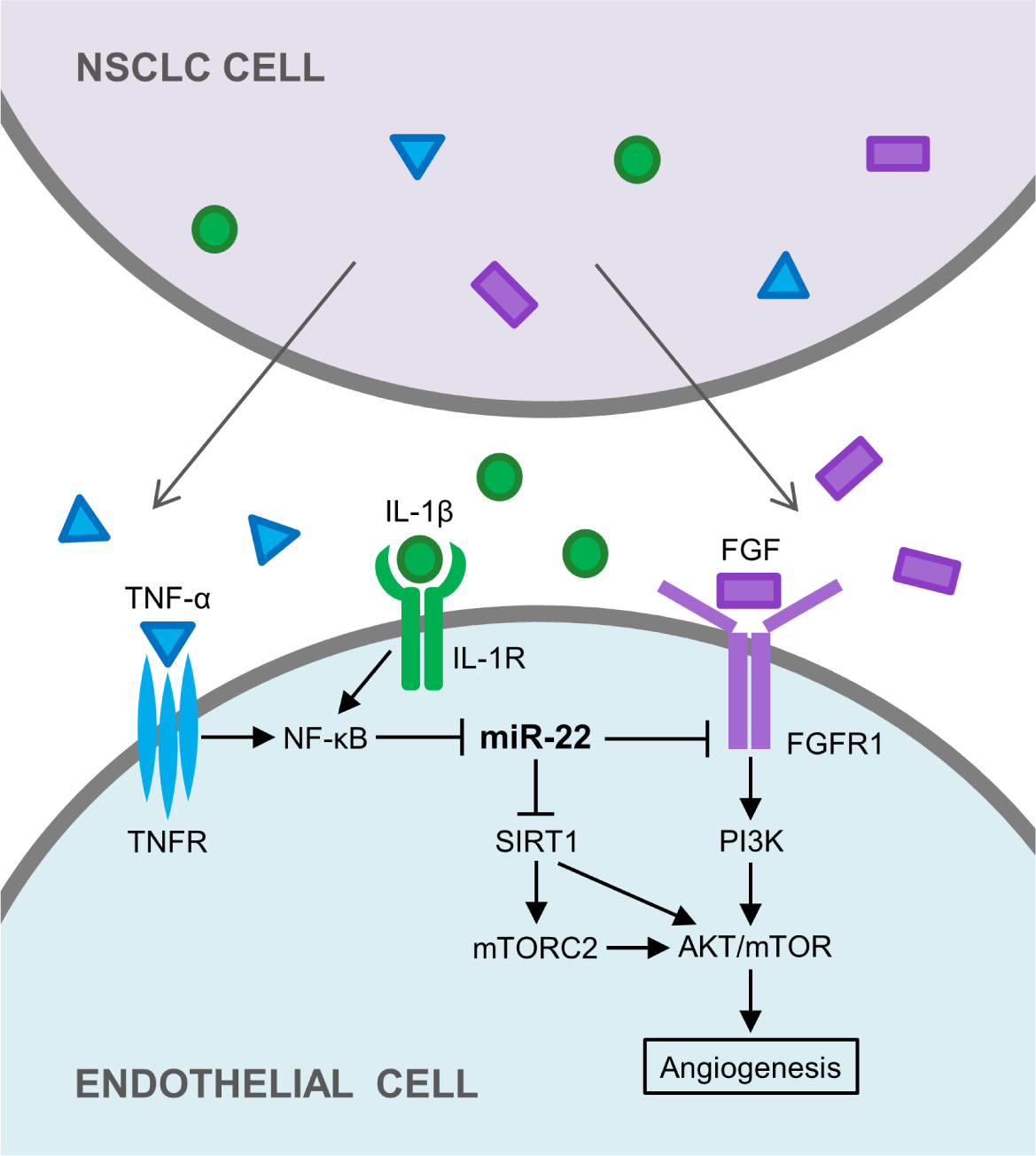
NSCLC cells induce angiogenesis by down-regulating endothelial miR-22, which targets *SIRT1* and *FGFR1*. The scheme summarizes the underlying mechanisms, as outlined in detail in the discussion section.

## Materials and Methods

### Study design

The main objective of our study was to analyze the function of endothelial miR-22 in regulating NSCLC angiogenesis. After identification of endothelial miR-22 to be significantly down-regulated in human NSCLC tissue from 12 patients, the following studies were designed: i) A contact and a non-contact coculture system were established *in vitro* using human ECs and NSCLC cells to study the regulation of endothelial miR-22 by NSCLC cells.

ii) A panel of *in vitro* assays were exploited to investigate the effects of miR-22 on major angiogenic steps, including EC proliferation, migration and tube formation. Mimic and inhibitor of miR-22 were transfected into ECs to perform gain- and loss-of-function studies.

iii) A Matrigel plug assay and a mouse flank tumor model were performed to confirm the *in vivo* inhibitory effects of miR-22 on angiogenesis and tumor growth. iv) Real-time PCR, Western blot and luciferase assays were used to identify and verify the target genes of miR-22. In this study, the sample size was estimated based on previous publications and experience. For each *in vitro* assay, at least 3 independent experiments with at least 3 biological replicates were performed to ensure the reproducibility and replicability of the results. Biological replicates are defined as separate cell cultures processed at the same time. Each mouse model included at least 5 mice in each group. These *in vivo* experiments could not be randomized, but all the analyses were performed by the investigators blinded to group assignment. All collected data were included in the analysis and no outliers were excluded.

### Chemicals

The NF-κB inhibitor Bay 11-7082, SIRT1 inhibitor EX-527 and FGFR1 inhibitor PD173074 were purchased from Santa Cruz Biotechnology (Heidelberg, Germany). The AKT inhibitor MK-2206 2HCL (MK-2206) was purchased from SelleckChem (Munich, Germany).

### Patient samples

Human NSCLC tissues and matched adjacent non-tumor lung tissues were obtained from 12 patients with lung adenocarcinoma. The pathological characteristics of these patients are shown in Supplementary file 1. All samples were dissected by professional pathologists in Saarland University Hospital, fixed in 4% formalin and embedded in paraffin. This study has been approved by the local ethics committee (permit number: 01/08) and the informed consent was provided by the patients.

### LCM

Sections with a thickness of 5 µm of lung adenocarcinoma and matched adjacent non-tumor lung tissue were mounted on MembraneSlides (Leica Microsystems, Wetzlar, Germany) and stained with hematoxylin and eosin. By using a microdissection microscope (Leica AS LMD, Leica), ECs were dissected and catapulted into the cap of 0.5 mL tubes (Leica) after removal of blood cells from capillaries. Approximately 2,000 ECs were retrieved from each sample. This procedure was assisted by an experienced pathologist.

### Cell culture

HDMECs (PromoCell, Heidelberg, Germany) were cultured in endothelial cell growth medium (EGM)-MV (PromoCell). HUVECs (PromoCell) were cultured in EGM (PromoCell). NHDFs (kind gift from Dr. Wolfgang Metzger, Department of Trauma, Hand and Reconstructive Surgery, Saarland University, Germany) were cultured in Dulbecco’s modified Eagle’s medium (DMEM; PAA, Cölbe, Germany) supplemented with 10% fetal calf serum (FCS), 100 U/mL penicillin and 0.1 mg/mL streptomycin (PAA). hPC-PLs (PromoCell) were cultured in pericyte growth medium (PromoCell). The human NSCLC cell lines NCI-H460 and NCI-H23 (ATCC, Wesel, Germany) were maintained in RPMI 1640 medium supplemented with 10% FCS, 100 U/mL penicillin and 0.1 mg/mL streptomycin. All cells were incubated at 37 °C in a humidified atmosphere containing 5% CO_2_.

### Cell co-culture

Contact and non-contact co-culture systems were used to assess the influence of tumor cells on endothelial miR-22 expression. For contact co-culture, 1×10^6^ HDMECs with or without 5×10^6^ NCI-H460 or NCI-H23 cells were seeded into 100-mm dishes and cultured in FCS-free endothelial cell basal medium (EBM) for 24 h. HDMECs were then isolated using a human CD31 MicroBead kit (Miltenyi Biotec, Bergisch Gladbach, Germany) according to the manufacturer’s instructions. Briefly, the co-cultured cells were detached with accutase (PAA) and suspended in 100 μL EBM. Subsequently, 30 µL FcR blocking reagent and 30 µL CD31 MicroBeads were added, followed by incubation at 4 °C for 15 min. After adding 1 mL EBM, the cells were sequentially collected by centrifugation, resuspended in 1 mL EBM and applied onto the LS Columns in the magnetic field of a MidiMACS separator. The column was washed 10 times with 3 mL EBM and then removed from the separator. The retained endothelial cells were flushed out 3 times with 4 mL EBM by pushing the plunger into the column and collected for further purity assessment and RNA extraction. For non-contact co-culture, 6-well transwell plates containing inserts with 0.4 μm pores (Corning, Wiesbaden, Germany) were used, which allowed soluble factors but not cells to pass through. A number of 1×10^5^ NSCLC cells were loaded onto the inserts and 2×10^5^ HDMECs were plated in the wells. After culture in EBM for 24 h, HDMECs were collected for RNA extraction.

### Immunocytochemistry

To check the cellular localization of p65, HDMECs were seeded on coverslips placed in a 6-well transwell plate and NCI-H460 cells were loaded onto the inserts. After culture in EBM for 4 h, HDMECs were fixed in 3.7% paraformaldehyde for 30 min, permeabilized with 0.5% Triton X-100 for 10 min and blocked with 2% bovine serum albumin (BSA) for 15 min. Afterwards, the cells were incubated with a primary antibody against p65 (1:25; R&D systems, Wiesbaden, Germany) for 1 h followed by the incubation with a Cy3-conjugated secondary antibody (1:250; Abcam, Cambridge, UK) for another 1 h. Cell nuclei were stained with Hoechst 33342 (Sigma-Aldrich, Taufkirchen, Germany). The percentage of p65-positive nuclei was quantified in 8 regions of interest (ROIs) of each coverslip at 40× magnification with a BX-60 microscope (Olympus, Hamburg, Germany).

### Cell transfection

To investigate the function of miR-22 in HDMECs, the cells were transfected with miR-22m (Qiagen, Hilden, Germany) or miR-22i (Qiagen) for 48 h to up- or down-regulate intracellular miR-22, respectively. Transfection reagent HiPerFect (Qiagen) was used according to the manufacturer’s protocol. Cells transfected with NCm (Qiagen) or NCi (Qiagen) served as controls.

### WST-1 assay

To assess cell viability, WST-1 assays (Roche Diagnostics, Mannheim, Germany) were performed according to the manufacturer’s instructions. Briefly, 4×10^3^ HDMECs were seeded in 96-well plates and incubated for the indicated time periods. Then, 10 µL WST-1 reagent was added into each well. After 30 min of incubation, the absorbance of each well was measured at 450 nm with 620 nm as reference by a microplate reader (PHOmo; anthos Mikrosysteme GmbH, Krefeld, Germany). The control group was assigned a value of 100%.

### Flow cytometry

To analyze the purity of isolated HDMECs, the cells were incubated with a fluorescein isothiocyanate (FITC)-conjugated mouse anti-human CD31 antibody (1:50; BD Pharmingen, San Diego, CA, USA) for 30 min at room temperature followed by 3 washes with phosphate buffered saline (PBS). At least 10,000 events were acquired using a FACScan flow cytometer (BD Biosciences, Heidelberg, Germany) and analyzed with CellQuest Pro software (BD Biosciences).

The function of miR-22 in cell cycle regulation was also detected by flow cytometry as previously described (31). Briefly, transfected HDMECs were reseeded and incubated for 24h. The cells were then collected and fixed followed by staining with propidium iodide (PI) and digestion with RNase A (Sigma-Aldrich). Subsequently, the cell cycle distribution was assessed by the FACScan flow cytometer and the DNA histograms of 10,000 cells were analyzed with the BD CellQuest Pro software.

### Cell migration assay

To evaluate EC motility, two different migration assays were performed. For the scratch wound healing assay, HDMECs were seeded in 35-mm culture dishes. After reaching confluence, the cell monolayer was scratched with a 10-µL pipette tip to generate scratch wounds and then rinsed with PBS to remove non-adherent cells. Phase-contrast microscopy (BZ-8000; Keyence, Osaka, Japan) was used to observe the wounds immediately after scratching (0 h) as well as after 12 h or 24 h. The wound area was measured and expressed as a percentage of corresponding NCm or NCi controls.

The transwell migration assay was performed as previously described (32). Briefly, 2.5×105 transfected HDMECs in 500 µL EBM were seeded into an insert of 24-transwell plates with 8 μm pores (Corning) and 750 µL EBM supplemented with 1% FCS was added to the lower well. Cells were allowed to migrate for 5 h and thereafter stained with Dade Diff-Quick (Dade Diagnostika GmbH, Munich, Germany). Cell migration was quantified by counting the number of migrated cells in 20 ROIs at 20× magnification using a BZ-8000 microscope (Keyence) and expressed as a percentage of corresponding NCm or NCi controls.

### Tube formation assay

To assess the tube forming activity of ECs, 1.5×10^4^ transfected HDMECs were added into each well of a 96-well plate pre-coated with 50 µL Matrigel (∼10 mg/mL; Corning). After incubation for 18 h, the formation of tubular structures was observed under phase-contrast microscopy (BZ-8000; Keyence). Tube formation was quantified by analyzing the number of meshes (i.e. areas completely surrounded by endothelial tubes) with the ImageJ software (U.S. National Institutes of Health, Bethesda, Maryland, USA) and expressed as a percentage of corresponding NCm or NCi controls.

### Aortic ring assay

To investigate the function of miR-22 in aortic sprouting, aortic rings processed from male BALB/c mice (8 weeks old) were transfected for 18 h with 50 nM miR-22m, 1 µM miR-22i or scrambled NCm and NCi, and then embedded in Matrigel (∼10 mg/mL; Corning) in a 96-well plate. After Matrigel polymerization, DMEM supplemented with 10% FCS was added into each well and sprouts from the aortic wall were allowed to develop for 6 days followed by observation with phase-contrast microscopy (BZ-8000; Keyence). Aortic sprouting was quantified by measuring the area of the outer aortic vessel sprouting and expressed as a percentage of corresponding NCm or NCi controls.

### Animal models

All animal experiments were approved by the local governmental animal protection committee (permit number: 22/2014) and were conducted in accordance with the German legislation on protection of animals and the NIH Guidelines for the Care and Use of Laboratory Animals (NIH Publication #85-23 Rev. 1985).

To investigate the *in vivo* function of miR-22 in angiogenesis, a Matrigel plug assay was performed as previously described (17). Briefly, transfected HDMECs in EBM (1×10^7^ cells/mL) were mixed with the same volume of growth factor-reduced Matrigel (∼20 mg/mL; Corning) and then supplemented with 1 µg/mL VEGF (R&D Systems), 1 µg/mL bFGF (R&D Systems) and 50 IU/mL heparin (B. Braun, Melsungen, Germany). Then, 300 µL Matrigel admixed with HDMECs was subcutaneously injected into 8-10-week-old CD1 nude mice (∼25 g). The Matrigel plugs were collected for immunohistochemical analyses 7 days after implantation.

The function of endothelial miR-22 in tumor angiogenesis and growth was evaluated in a flank tumor model. For this purpose, 1.5×10^5^ NCI-H460 cells in combination with 1.5×10^6^ NCm- or miR-22m-transfected HDMECs were suspended in 50 µL EGM-MV and injected subcutaneously into the flanks of 8-week-old NOD-SCID (NOD. CB17/AIhnRj-Prkdc^scid^) mice (Janvier Labs, Le Genest-St-Isle, France). Two perpendicular diameters of the developing tumors were repetitively measured on day 0, 3, 7, 10 and 14 by means of a caliper. The tumor volumes were calculated using the formula V = 1/2 (L × W^2^), where L was the longer and W was the shorter diameter (33). The tumor development was also assessed using a combined ultrasound and photoacoustic imaging system (Vevo LAZR) with a LZ550 scanhead (40 MHz center frequency) (FUJIFILM VisualSonics Inc, Toronto, Canada) on day 10 and 14 after implantation. The ultrasound images of tumors were analyzed by means of a three-dimensional reconstruction using VisualSonics software (Vevo LAB 1.7.2.). At the end of the experiment, i.e. on day 14, the tumors were carefully excised, weighed and further processed for immunohistochemical analyses.

### Immunohistochemistry

Formalin-fixed specimens of Matrigel plugs and tumors were embedded in paraffin and 2-µm sections were cut. To detect the neovascularization of the plugs and tumors, the sections were stained with a rabbit anti-human CD31 antibody (1:100; Abcam) or a rabbit anti-mouse CD31 antibody (1:100; Abcam), followed by a goat-anti-rabbit Alexa Fluor 555-labeled secondary antibody (1:100; Life Technologies, Eugene, OR, USA) or a goat-anti-rat Alexa Fluor 488-labeled secondary antibody (1:100; Life Technologies). Cell nuclei were stained with Hoechst 33342 (Sigma-Aldrich). The sections were subsequently examined using a fluorescence microscope (BX60; Olympus). Microvessel density was quantified by counting the numbers of CD31-positive microvessels in 10 ROIs of each section at 20× magnification. To evaluate the proliferation and apoptosis of tumor cells, sections were stained with a monoclonal rabbit antibody against Ki67 (1:400; Cell Signaling Technology, Frankfurt, Germany) or a polyclonal rabbit antibody against cleaved casp-3 (1:100; New England Biolabs, Frankfurt, Germany), followed by a biotinylated goat anti-rabbit secondary antibody (Abcam) and streptavidin-peroxidase conjugate (ready-to-use; Abcam). The staining was completed by incubation with 3-amino-9-ethylcarbazole substrate (Abcam) before the sections were counterstained with Mayers hemalaun solution (HX948000; Merck, Darmstadt, Germany). The percentages of Ki67-positive proliferating and cleaved casp-3-positive apoptotic tumor cells were quantified in 12 ROIs of each section at 40× magnification with a BX-60 microscope (Olympus).

### Quantitative real-time polymerase chain reaction (PCR)

Total RNA was extracted using RNeasy FFPE Kit (Qiagen), RNeasy Mini kit (Qiagen) or miRNeasy Mini kit (Qiagen) following the manufacturer’s instructions. Then, the extracted RNA was processed for the reverse transcription reaction by utilizing QuantiTect Reverse Transcription Kit (Qiagen) or miScript II RT Kit (Qiagen). Noteworthy, after reverse transcription, cDNA of dissected ECs by LCM was further amplified using miScript PreAMP PCR Kit (Qiagen). Quantitative real-time PCR was performed and analyzed in a MiniOpticon Real-Time PCR System (BioRad, Munich, Germany) using QuantiTect SYBR green PCR Kit (Qiagen) or miScript SYBR Green PCR Kit. The relative expression levels of genes and miRNAs were calculated using the 2^-ΔΔCt^ method with GAPDH and U6 as endogenous control, respectively. Gene-specific primer sequences are listed in Supplementary file 2. To analyze mature miRNA expression, miScript primer assays for Hs_miR-22_1 and Hs_RNU6-2_11 from Qiagen were used.

### Western blot analysis

As previously described (34), whole cell lysates were separated on 8% sodium dodecyl sulfate (SDS) polyacrylamide gels and transferred to polyvinylidene difluoride (PVDF) membranes (BioRad). The membranes were blocked and incubated overnight at 4 °C with a mouse monoclonal anti-FGFR1 antibody (1:100; Cell Signaling Technology), a rabbit polyclonal anti-p-AKT antibody (1:500; Cell Signaling Technology), a rabbit monoclonal anti-AKT antibody (1:500; Cell Signaling Technology), a rabbit monoclonal anti-p-mTOR (1:500; Cell Signaling Technology), a rabbit monoclonal anti-mTOR (1:500; Cell Signaling Technology) or a mouse monoclonal anti-β-actin antibody (1:2,000; Sigma-Aldrich). This was followed by the corresponding horseradish peroxidase (HRP)-conjugated secondary antibodies (1:3,000; GE Healthcare, Freiburg, Germany). An electrochemiluminescence assay (GE Healthcare) was then performed and signals were acquired using a ChemoCam Imager (Intas, Göttingen, Germany). The intensities of protein bands were analyzed using the ImageJ software (U.S. National Institutes of Health).

### Luciferase assay

For target validation, a control luciferase reporter plasmid (CmiT000001-MT06; GeneCopoeia, Rockville, USA) or *FGFR1*-3’UTR target plasmid (HmiT005432-MT06; GeneCopoeia) was co-transfected with 50 nM NCm or miR-22m into 293T cells using Lipofectamine 2000 (Invitrogen). After 48 h of incubation, Renilla and Firefly luciferase activities were measured by the Dual-Luciferase Reporter Assay Kit 2.0 (GeneCopoeia) using a Tecan Infinite 200 Pro microplate reader (Tecan, Crailsheim, Germany). Relative luciferase activity was quantified by normalizing the Firefly luciferase signal to that of Renilla luciferase and expressed as a percentage of NCm controls.

## Statistics

Statistical comparisons between two groups were made by the paired Student’s t-test (for the analysis of patient samples) or the unpaired Student’s t-test using GraphPad Prism 9. Statistical comparisons between multiple groups were made by one-way ANOVA followed by the Tukey’s multiple comparisons test using GraphPad Prism 9. All data were expressed as means ± SEM. A value of P<0.05 was considered significant.

## Acknowledgments

We are grateful for the excellent technical assistance of Janine Becker, Christina Max and Ruth Nickels (all from the Institute for Clinical and Experimental Surgery, Saarland University). This work was supported by a research grant to Y.G. from the Medical Faculty of Saarland University (HOMFORexzellent 2015).

## Figure legends for figure supplement

**Figure 2-figure supplement 1.**
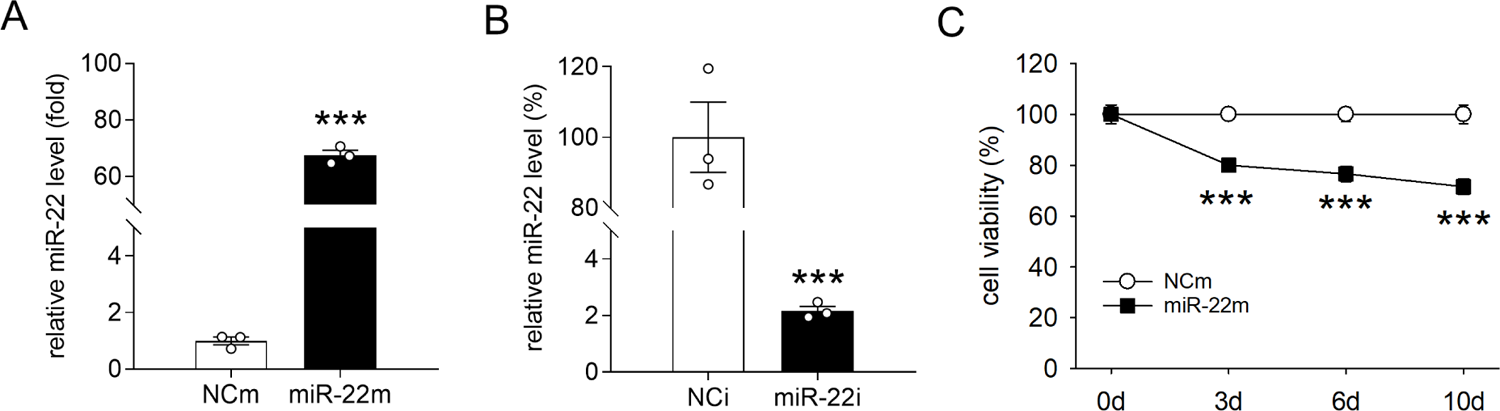
Expression of miR-22 and its effect on EC viability. **A**, **B**: Expression level of miR-22 (in fold of NCm or in % of NCi) in HDMECs transfected with miR-22m (**A**), miR-22i (**B**) or corresponding scrambled NCm (**A**) and NCi (**B**), as assessed by real-time PCR (n = 3). **C**: Viability (in % of NCm) of HDMECs transfected with miR-22m or NCm, as assessed by WST-1 assay (n = 4). After transfection, the cells were reseeded in 96-well plates and cultured for 0, 3, 6 or 10 days followed by WST-1 assay. Means ± SEM. ***P < 0.001 vs. NCm or NCi.

**Figure 2-figure supplement 2.**
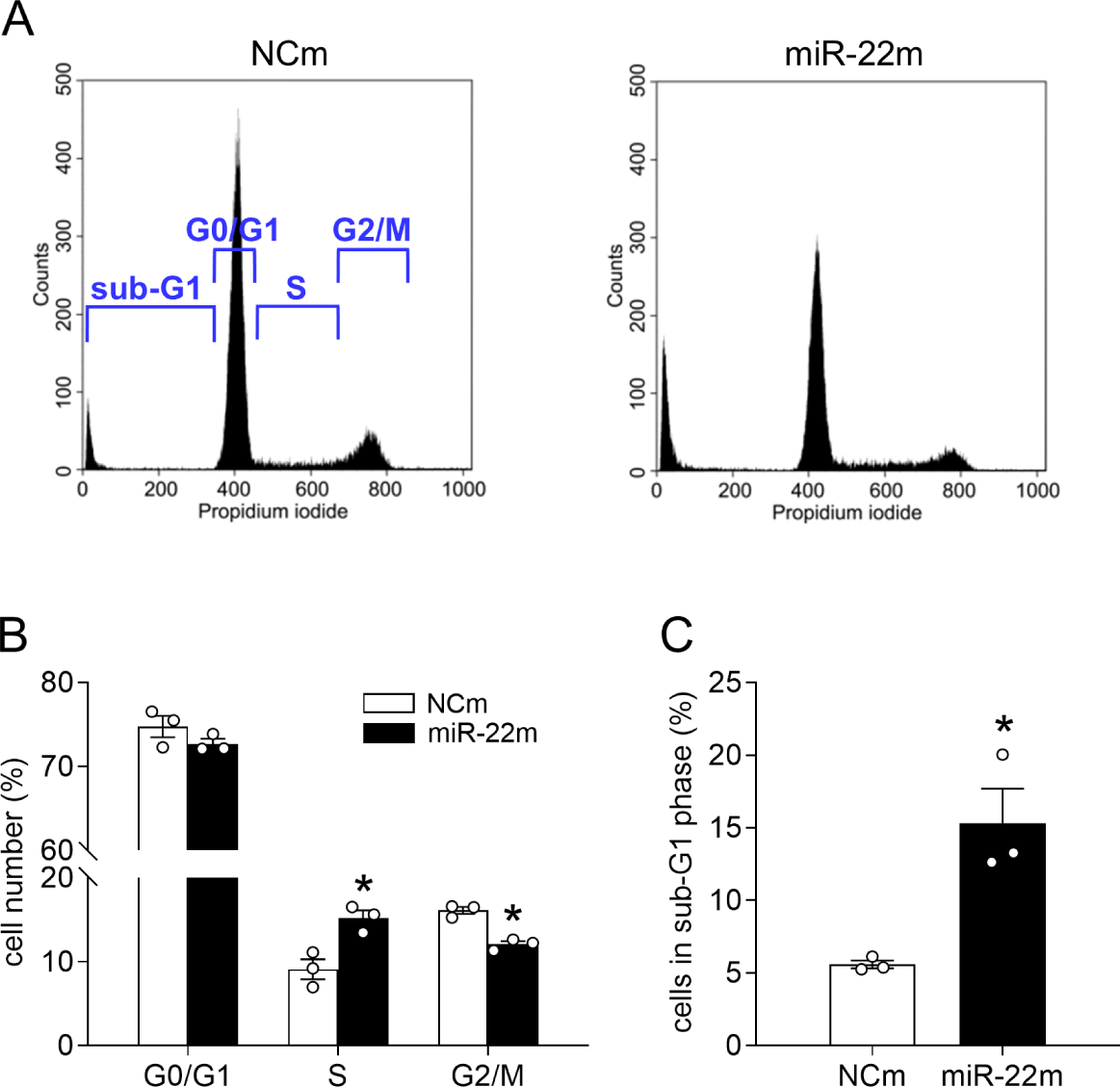
MiR-22 blocks HDMECs in the S phase. **A**: Representative cell cycle analysis of HDMECs transfected with NCm or miR-22m, as assessed by flow cytometry. **B**: Number of NCm- or miR-22m-transfected HDMECs in the G0/G1, S and G2/M phase (in % of total cell number), as assessed by flow cytometry (n = 3). **C**: Number of NCm- or miR-22m-transfected HDMECs in the sub-G1 phase (in % of total cell number), as assessed by flow cytometry (n = 3). Means ± SEM. *P < 0.05 vs. NCm.

**Figure 2-figure supplement 3.**
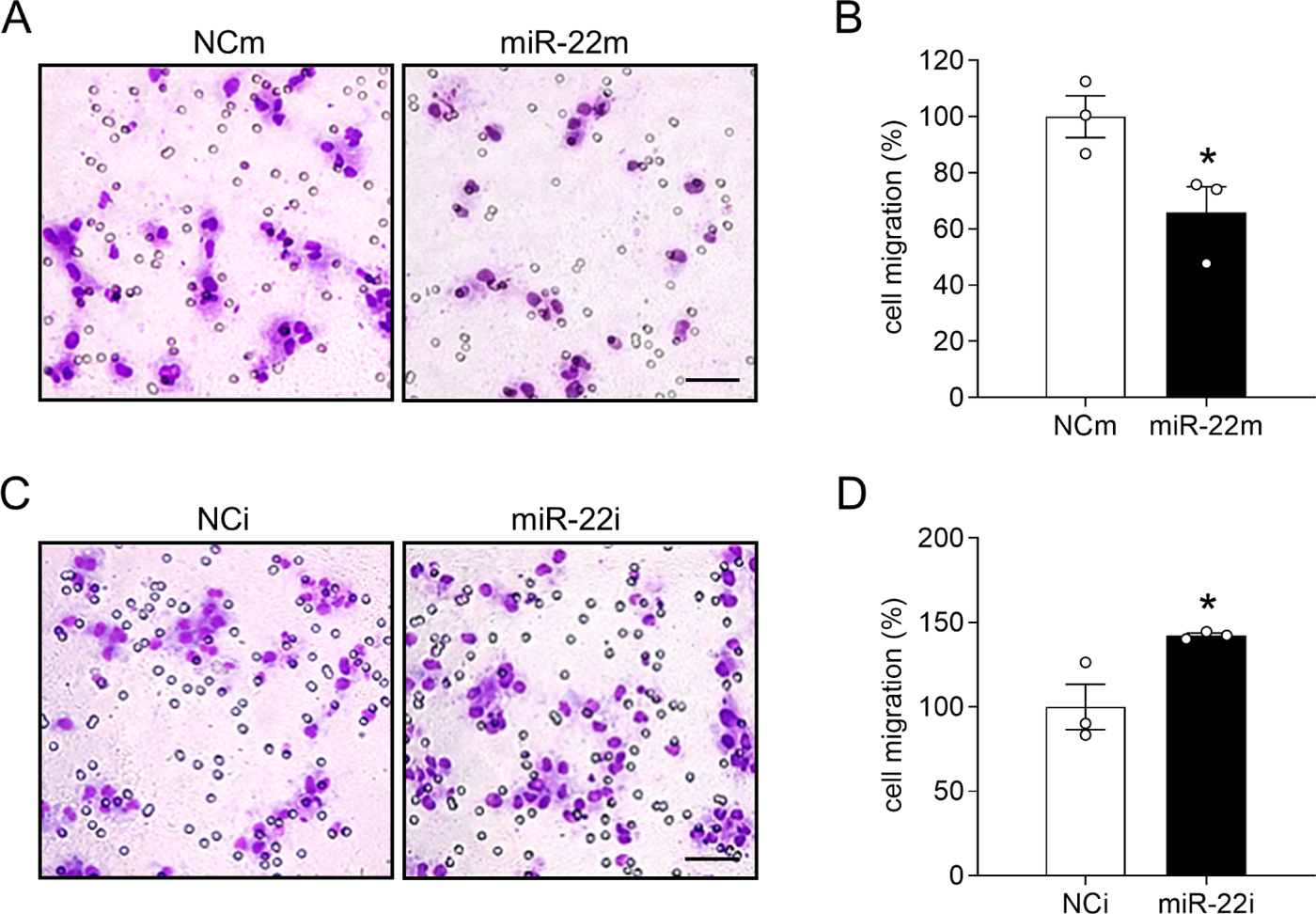
MiR-22 suppresses HDMEC migration. **A**, **C**: Light microscopic images of migrated HDMECs. The cells were transfected with miR-22m (**A**), miR-22i (**C**) or corresponding scrambled NCm (**A**) and NCi (**C**). Scale bars: 55 µm. **B**, **D**: Migration (in % of NCm or NCi) of HDMECs transfected with miR-22m (**B**), miR-22i (**D**) or corresponding scrambled NCm (**B**) and NCi (**D**), as assessed by transwell migration assay (n = 3). Means ± SEM. *P < 0.05 vs. NCm or NCi.

